# GFP-Tel1 is a marker of reversed replication forks that reveals constant fork backtracking during deoxyribonucleotide imbalance

**DOI:** 10.64898/2026.11.12.766229

**Authors:** Morgane Auboiron, Kseniya Samardak, Antoine Barthe, Philippe Pasero, Benjamin Pardo, María Moriel-Carretero

**Affiliations:** Centre de Recherche en Biologie cellulaire de Montpellier (CRBM), Université de Montpellier - Centre National de la Recherche Scientifique, Montpellier, France; Institut de Génétique Humaine (IGH), Université de Montpellier - Centre National de la Recherche Scientifique, Montpellier, France

**Keywords:** Tel1, replicative stress, fork reversal, nucleoplasmic foci

## Abstract

The apical kinase of the DNA Damage Response Tel1/ATM participates in at least three transactions key to the preservation of genome integrity: maintenance of a homeostatic telomere length, signaling of double strand breaks (DSB) in the DNA and a yet-poorly-defined contribution at reversed forks, a structure arising during problems in DNA replication. In a previous work, by tagging Tel1 in its N-terminus with yeast Enhanced Green Fluorescent Protein, we discovered that Tel1 forms foci in close proximity to the nuclear periphery in response to both chemically and enzymatically induced DSB. Here, we explore the determinants of this localization to the nuclear envelope, and we uncover that the subnuclear localization of the foci are a discriminant feature of Tel1’s roles: foci forming in the interior of the nucleus mark reversed replication forks. In contrast, Tel1’s function at telomeres are not tractable using GFP-Tel1 as a tool. Importantly, and in contrast to the prevalent notion that fork reversal is a phenomenon greatly restricted to metazoan replication forks, our characterization in the yeast *S. cerevisiae* of GFP-Tel1 at reversed forks allows us to propose a reconciling principle for how fork reversal contributes to tolerating replicative stress across evolutionary distant organisms.

## Introduction

The integrity of the genetic material is continuously surveilled to ensure organism fitness. This way, cells confronted to genotoxic challenge activate the DNA damage response (DDR), a process that concurs with detection of the problem by dedicated sensors, and the subsequent engagement of upstream kinases. Mainly two members of the phosphoinositide 3-kinase-related kinases (PIKK) family, namely Tel1/ATM and Mec1/ATR (as known in *Saccharomyces cerevisiae*/humans), take care of this task. These kinases launch the phosphorylation of downstream mediator and effector proteins, among which Rad53/CHK1-CHK2, in charge of launching DNA repair and other protective strategies (Blackford & Jackson, 2017). The DDR does not display a unique program of action, but accommodates, with different deployed profiles, to the type of DNA challenge. This way, depending, for example, on whether the DNA suffers from a DNA double strand break (DSB) or a replication fork that cannot continue duplicating DNA (a situation generically defined as replication stress), different responses will be engaged (Pardo *et al*, 2016).

The process of replication, accomplished by a large number of simultaneously progressing replisomes, is challenged whenever these have to cope with DNA lesions, tightly bound protein complexes, secondary DNA structures, programmed replication pause sites and natural activities occurring on DNA such as transcription (Aguilera & García-Muse, 2013; Zeman & Cimprich, 2014). Among the strategies implemented in front of these replicative challenges, forks can undergo reversal. Fork reversal implies the transition from a typical replication fork (which is a three-way junction molecule) into a four-way one. This occurs by the coordinated annealing of the two freshly synthesized strands and the re-annealing of the parental strands, the whole forming a backtracked structure with respect to the fork elongation point (Neelsen & Lopes, 2015). While fork reversal is now considered a common response in mammals to various types of DNA replication stresses, it is a phenomenon rarely detected in *S. cerevisiae* and, as such, considered as a by-product of replication failure in most scenarios (Neelsen & Lopes, 2015).

We have reported that the apical PIKK kinase Tel1, tagged in its N-terminus with yeast Enhanced Green Fluorescent Protein (yEGFP, for simplicity, GFP) forms foci in close proximity to the nuclear periphery in response to DSB, irrespective of these being induced by chemical genotoxins (zeocin), or enzymatically (Cas9) (Auboiron *et al*, 2025). These foci form soon after DSB occurrence, disappear fast once the source of DSB is removed, and precede the apparition of foci formed by the DNA repair factor Rad52, altogether suggesting that GFP-Tel1 foci mark early steps of DSB processing, i.e., DSB detection (Auboiron *et al*, 2025). Also, we showed that a single nucleus can display several GFP-Tel1 foci, with, in rare cases, up to eight of them. Given that the number of induced DSB upon zeocin treatment greatly exceeded eight, these data implied that one single GFP-Tel1 focus may gather several DSBs, which would be signaled simultaneously (Auboiron *et al*, 2025). To broaden our understanding of GFP-Tel1 foci formation and further explore its possibilities for discovery, in this work we have asked if GFP-Tel1 foci require Tel1’s kinase activity and whether other DNA substrates at which Tel1 has known functions, like telomeres and reversed forks, also congregate GFP-Tel1 foci.

We discover that the participation of Tel1 to different genome integrity preservation pathways can be tracked and discriminated by its ability to form nuclear foci of different positions. In more detail, we show that peripheral GFP-Tel1 foci mark the early stages of DSB recognition before Tel1 engages in any kinase activity, in contrast to foci forming in the interior of the nucleus, which mark reversed replication forks. However, the functions Tel1 deploys at telomeres are not tractable using GFP-Tel1 as a tool, likely because a minimum number of DNA substrates need to gather in space to give rise to a focus. Beyond, our establishment of this strategy, likely the first ever capable of monitoring reversed forks in *S. cerevisiae*, has allowed us to extract a reconciling principle for how fork reversal contributes to tolerating replicative stress across evolutionary distant organisms. As such, our data prompt to dismiss the duality claiming that reversed forks in response to replicative stress are an exclusive hallmark of metazoan cells that do not occur in yeast, and instead support that fork reversal occurs as a continuum of variable duration in all organisms.

## Results

### 1. GFP-Tel1 foci form in response to DNA double strand breaks irrespective of Tel1’s ability to phosphorylate its downstream targets

GFP-Tel1 foci formation occurs after several GFP-Tel1 molecules cluster together in space, reaching a fluorescence threshold that can be imaged. In agreement, the number of DSBs to be signaled exceeds the number of detected foci, further suggesting that several DNA breaks are being signaled in proximity (Auboiron *et al*, 2025). We however ignore if Tel1’s kinase activity on target substrates is important to allow this congregation or if, on the contrary, DNA-Tel1 nucleation precedes such phosphorylations. The same goes for the peripheral position of the foci, which, we reported, form in tight proximity to the nuclear envelope (Auboiron *et al*, 2025). To assess these questions, we took advantage of a deeply characterized point mutant in the C-terminal FATC domain (Figure 1A), *tel1-21*, that has been shown to *1)* encode a protein as stable as the wild-type one (Figure 1B); *2)* be recruited to chromatin in response to DSB as efficiently as the control; *3)* preserve its self-phosphorylation; *4)* be, however, incapable of correctly phosphorylating its substrates at this location (Figure 1C), (Ogi *et al*, 2015). We induced DSBs with zeocin in WT (wild type) and *tel1-21* cells, both bearing the GFP tag at an identical N-terminal position, and monitored the kinetics and the level of GFP-Tel1 foci formation. *tel1-21* mutant cells formed GFP-Tel1 foci that were of similar aspect to those formed by the WT (Figure 1D), accumulated with identical kinetics, yet double in frequency (Figure 1E). The mean number of foci at any timepoint was higher in the mutant than in the WT, and the mean cumulative value in the mutant greatly exceeded that in the WT (Figure 1F). We interpret that this is likely because new DSBs arise and add up to the foci in which signaling has been initiated, yet has not adequately progressed. Last, about 75% of the foci were close to the nuclear periphery, as frequently as we have observed in the WT (Figure 1G, see Material & Methods for the zoning strategy).

**Figure 1.**
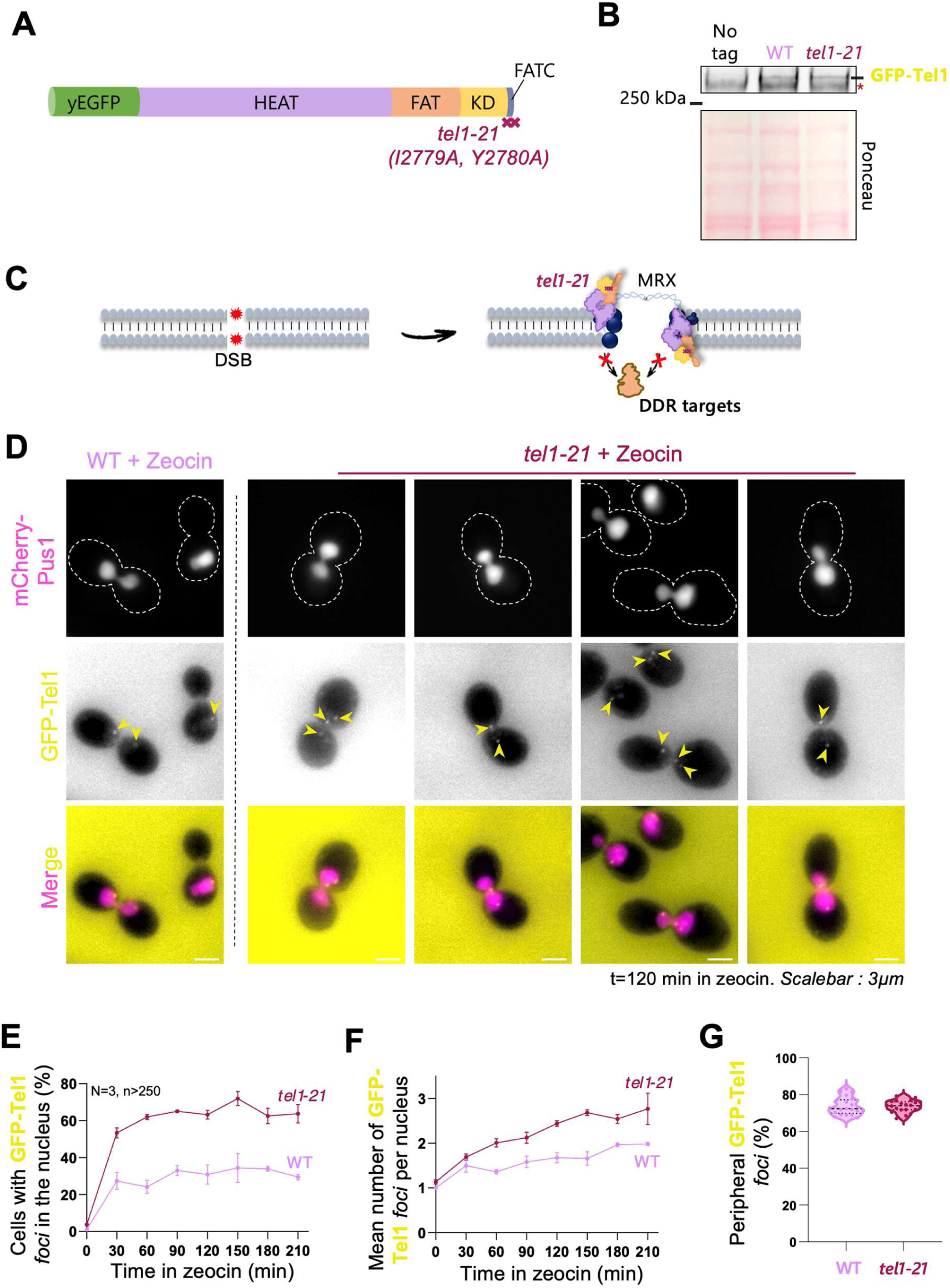
Tel1’s ability to phosphorylate its downstream targets is dispensable to form foci in response to DSB. **A**. Schematic representation of the *tel1-21* mutant tagged at the N-terminus with the yEGFP fluorophore. **B.** Western blot analysis of GFP-Tel1 in WT and *tel1-21* mutant strains. The red asterisk indicates a nonspecific band detected by the anti-GFP antibody. **C.** Schematic representation of *tel1-21* recruitment by the MRX complex in response to a DSB. Although recruited to DNA damage sites, this mutant is unable to phosphorylate downstream targets of the DNA damage response (indicated by crossed-out arrows). **D.** Representative fluorescence microscopy images of WT and *tel1-21* mutant cells after 120 min of 100 µg/mL zeocin treatment. Cell boundaries are outlined with dashed white lines. mCherry-Pus1 is used as a nuclear marker. GFP-Tel1 foci are indicated by yellow arrowheads. The white scale bar represents 3 µm. **E.** Quantification of the percentage of cells containing at least one nuclear GFP-Tel1 focus at each time point following 100 µg/mL zeocin treatment in WT and *tel-21* mutant strains. Each point is the mean of 3 independent experiments (N) considering at least 250 nuclei (n) per timepoint and experiment. The error bars represent the Standard Error of the Mean (SEM) out of the 3 independent experiments. **F.** Quantification of the mean number of GFP-Tel1 foci per positive nucleus in each strain. The total number of foci was divided by the number of positive nuclei. Details for reproducibility and plotting as in (E). **G.** Percentage of GFP-Tel1 foci localized at the nuclear periphery relative to nucleoplasmic foci. Each dot represents the percentage measured at a single time point from one biological replicate (7 timepoints across 3 independent biological replicates).

Together, we conclude that the phosphorylation of Tel1 targets after its recruitment at DSB is not necessary to permit its congregation on DNA in the form of foci, nor to warrant the reported peripheral position of the foci. More on the contrary, the inability to make signaling progress adequately likely delays the resolution of GFP-Tel1 foci, which accumulate in time and in space.

### 2. Telomeres do not support the formation of GFP-Tel1 foci

Telomeres, the end of chromosomes, represent another Tel1-binding substrate under given circumstances. Bound by the Rap1-Rif1-Rif2 complex when of normal length, the progressive loss of this protective complex at telomeres as they shorten with each replication cycle allows the binding of the Mre11-Rad50-Xrs2 (MRX) complex, which in turns recruits Tel1. Tel1 stimulates the activation of MRX nuclease function, which exposes a naked 3’ end, which in turn becomes a binding site for the Cdc13-Stn1-Ten1 (CST) complex, responsible for telomerase recruitment and telomere extension (reviewed in Shore & Bianchi, 2009 ; Figure 2A - left). Additionally, the very tip of telomeres, a protruding 3’ end, is also protected by the CST complex, whose binding to single stranded DNA is reminiscent of that displayed by Replication protein A (RPA). In fact, defects in CST binding permit RPA access to telomeric protruding 3’ ends, and subsequent stimulation of the other DDR kinase, Mec1.

**Figure 2.**
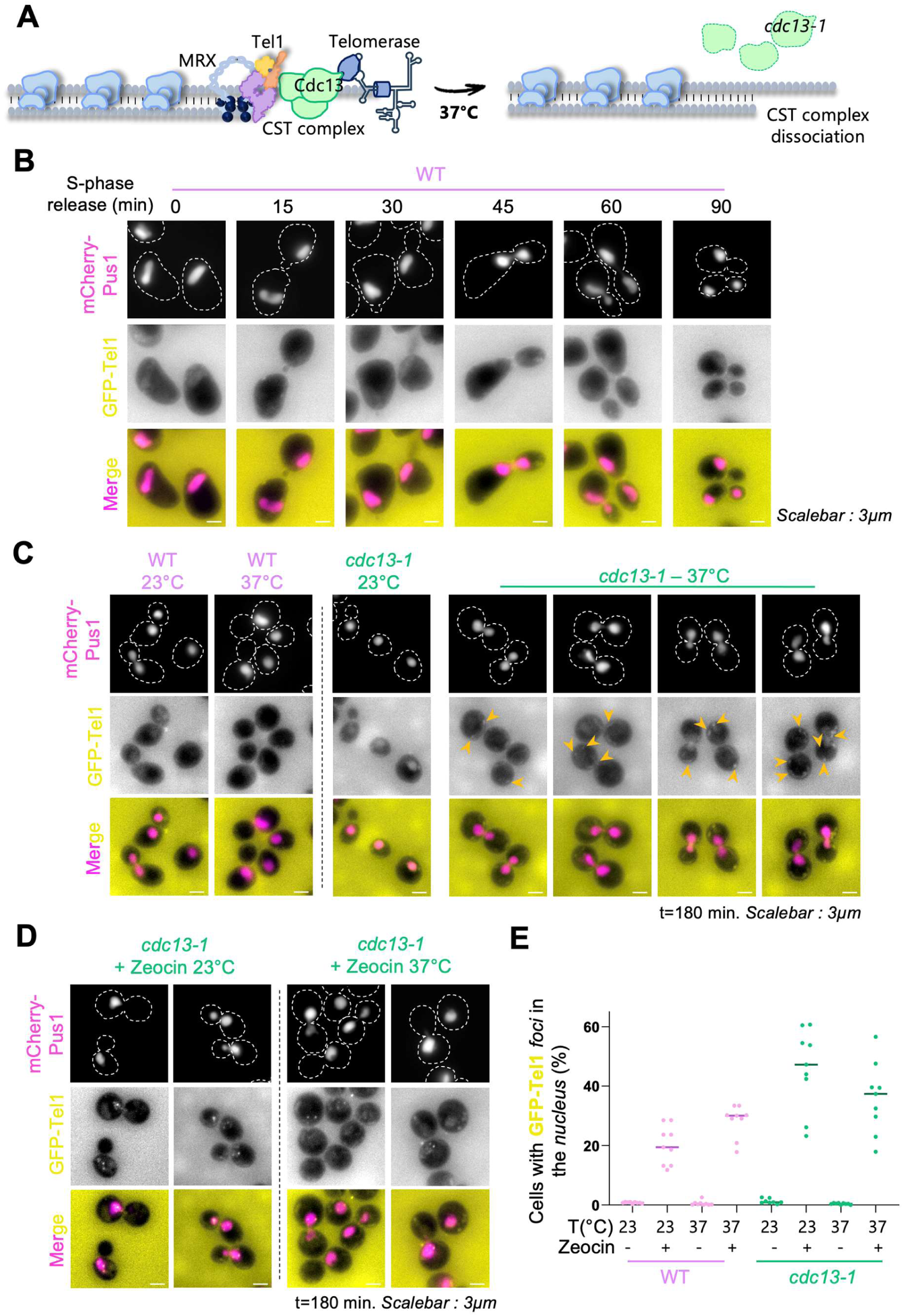
Telomeres do not support the formation of GFP-Tel1 foci. **A**. Schematic representation of telomere protection in WT cells, where the MRX complex recruits Tel1, which stimulates its nuclease activity, leading to the binding of the CST complex, itself recruiting and regulating telomerase activity for telomere extension. In the *cdc13-1* mutant, incubation at the restrictive temperature (37°C) leads to loss of Cdc13 function and telomere deprotection. **B.** Representative fluorescence microscopy images of WT cells following G_1_ synchronization and release into S phase. No GFP-Tel1 foci are detected at any time point under these conditions. **C.** Representative fluorescence microscopy images of WT and *cdc13-1* mutant cells after 180 min of incubation at permissive (23°C) or restrictive (37°C) temperatures. No nuclear GFP-Tel1 signal is detected at any time point under these conditions. Orange arrows point at cytoplasmic GFP-Tel1 signals observed in the *cdc13-1* mutant at the restrictive temperature. **D.** Representative fluorescence microscopy images of WT and *cdc13-1* mutant cells after 180 min of 100 µg/mL zeocin treatment. **For panels B-D.** Cell boundaries are outlined with dashed white lines. mCherry-Pus1 is used as a nuclear marker. The white scale bar represents 3 µm. **E.** Quantification of the percentage of cells displaying nuclear GFP-Tel1 localization under permissive or restrictive temperature conditions, with or without zeocin treatment. Each dot represents the percentage of positive nuclei measured at a single time point from one biological replicate (3 timepoints across 3 independent biological replicates). The colored horizontal bars indicate the mean of individual means.

We first interrogated if congregation of GFP-Tel1 foci could be seen in replicating WT cells that would mark naturally-occurring short-telomere detection events. To this end, we synchronized cells in the G_1_ phase of the cell cycle using α-factor and released them synchronously in replication (S) phase, monitoring at different intervals since release. Out of three independent experiments and at least 250 cells analyzed per timepoint and experiment, we failed to detect GFP-Tel1 foci in the cells at any of the acquired frames (Figure 2B). This suggested that either there are too few events occurring in the population and / or that the minimal threshold of Tel1 molecules that must gather in order to be detected as a focus was not reached in WT cells.

Although the presence of Tel1 is mostly documented for telomeres undergoing natural shortening, telomere deprotection, mostly known to stimulate the activation of the kinase Mec1 (see above), also requests Tel1 participation. In this sense, Tel1 limits the early accumulation of Exo1-dependent telomeric ssDNA when telomeres become deprotected, and its absence exacerbates the temperature sensitivity of *cdc13-1* cells (Zhang *et al*, 2018). To enquire if Tel1 can form foci in this pathological context, we used a thermosensitive mutant of the Cdc13 subunit of the CST complex, *cdc13-1* (Garvik *et al*, 1995; Weinert & Hartwell, 1993), in which we can impose a sudden telomere deprotection when rising the culturing temperature from 23°C to 37°C (Figure 2A). When cultured at the restrictive temperature (37°C), the *cdc13-1* mutant did not show any GFP-Tel1 foci formation (Figure 2C). This lack of foci formation was not due to the mutant being incapable of forming foci, as challenging the cells with zeocin-induced DSB led to the formation of foci with even higher frequency than the WT cells, and this irrespective of the temperature (Figure 2D,E).

It is worth commenting that frequent, bright GFP-Tel1-positive structures were observed out from the nucleus in this mutant when cultured at the restrictive temperature even in the case in which Tel1 is actively recruited to the nucleus in response to DSB induction with zeocin (Figure 2C orange arrows, Figure 2D - see discussion). In sum, we conclude that normal or deprotected telomeres do not support the formation of GFP-Tel1 foci.

### 3. GFP-Tel1 foci allow the distinction between reversed replication forks and DSBs by their differential nuclear localization

The last structure reported to date to attract Tel1 also resembles a DSB and is formed during the process of DNA replication when the two newly synthesized strands anneal with each other, forcing the parental strands to zip back together. In this process, termed fork reversal, the replication fork backtracks, which gives rise to a DNA end that mimics a DSB, at the tip of newly synthesized strands (Figure 3A). Tel1 recruitment to this tip has been reported to repel the engagement of the nuclease Mre11, which otherwise would degrade the extruded DNA (Menin *et al*, 2018). Fork reversal is reported as a physiological response to obstacles preventing the progression of the replication fork and has been observed by electron microscopy (EM) after psoralen cross-linking in response to various genotoxic treatments in human cells (Zellweger *et al*, 2015). In the yeast *S. cerevisiae*, fork reversal was formally demonstrated by EM in response to topoisomerase I trapping with camptothecin (CPT) (Menin *et al*, 2018). We wished to ask if GFP-Tel1 could also form foci in response to CPT-induced fork reversal, for which we exposed otherwise WT cells growing asynchronously to 100 µM CPT, then monitored them in time. CPT treatment elicited the formation of GFP-Tel1 foci (Figure 3B) in about 17% of the cells, a value that was mostly maintained in time (Figure 3C). To make sure that these foci represented *bona fide* reversed forks, we repeated the experiments in cells lacking the protein Mrc1 (*mrc1Δ*), an actor supporting fork progression proficiency and whose presence is necessary to permit CPT-induced fork reversal in yeast cells (Menin *et al*, 2018). Mrc1 absence prevented the formation of GFP-Tel1 foci induced by CPT (Figure 3C), arguing in favor of these foci most probably marking the tip of reversed forks. An important morphological difference of our analysis was that the foci we ascribed as stemming from reversed forks were very frequently occupying a nuclear central position (at about 65% of the foci-containing cells), named “nucleoplasmic foci” from here on (Figure 3D), in striking contrast with the foci associated to DSB, which are mostly peripheral (Figure 1, 75% of foci ; Auboiron *et al*, 2025). Thus, GFP-Tel1 appears to mark reversed forks as foci that form in the interior of the nucleoplasmic space.

**Figure 3.**
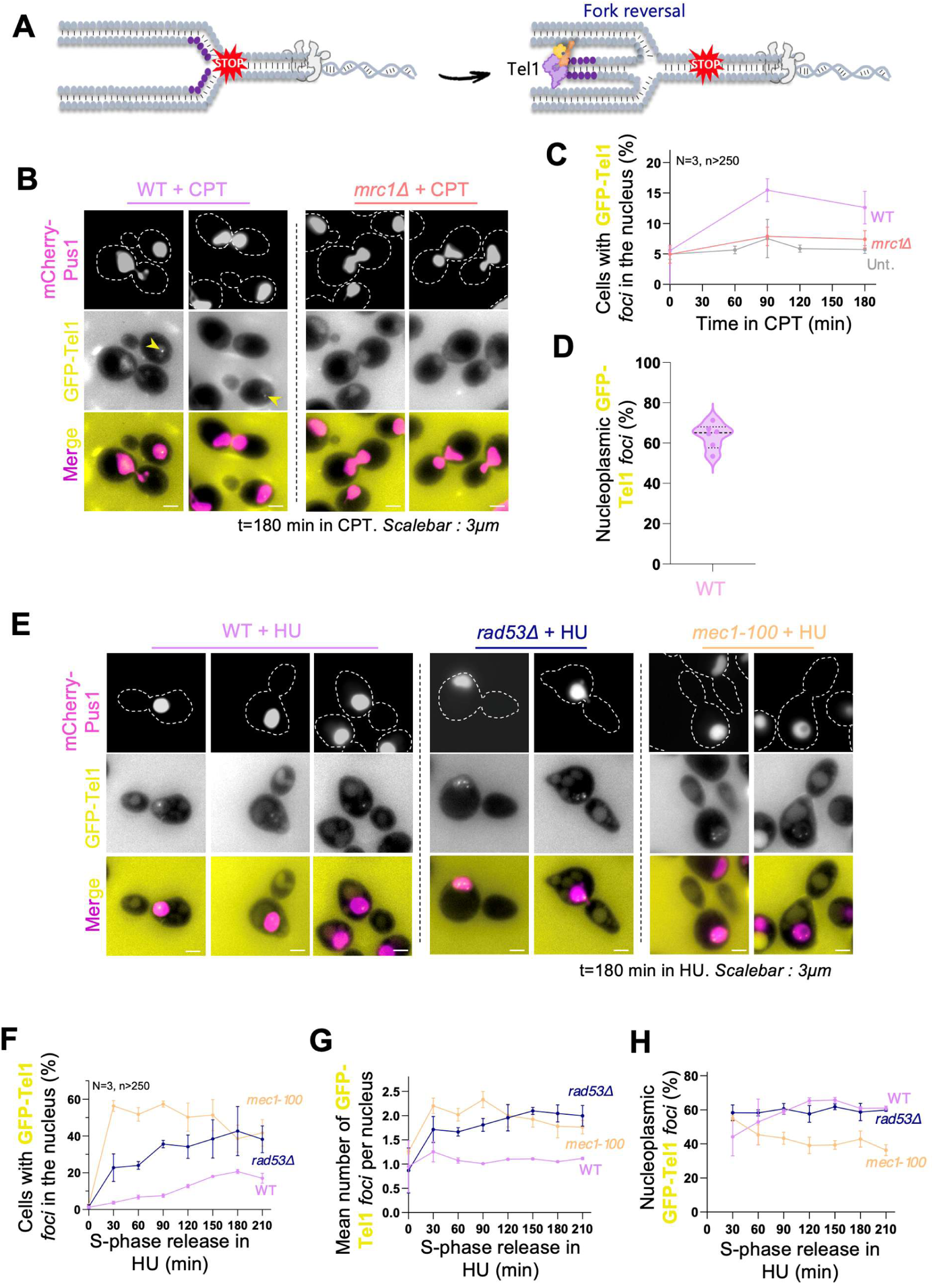
GFP-Tel1 foci allow to distinguish reversed replication forks from DSBs by their differential nuclear position. **A**. Schematic representation of Tel1 recruitment to the tip of a reversed replication fork. The same deoxyribonucleotides, shown in purple, are depicted before and after replication fork reversal to illustrate their displacement during the process. **B.** Representative fluorescence microscopy images of WT and *mrc1Δ* strains after 180 min of CPT treatment (100 µM final concentration). **C.** Quantification of the percentage of cells containing at least one nuclear GFP-Tel1 focus at each time point following 100 µM CPT treatment in WT and *mrc1Δ* strains. “Unt.” corresponds to untreated WT cells used as a control. **D.** Percentage of GFP-Tel1 foci localized in the nucleoplasm relative to foci localized at the nuclear periphery. Each dot represents the percentage measured at a single time point from one biological replicate (2 timepoints across 3 independent biological replicates). **E.** Representative fluorescence microscopy images of WT, *rad53Δ*, and *mec1-100* strains following G_1_ synchronization and 180 min after release into S phase in the presence of 200 mM HU. **For panels B and E.** Cell boundaries are outlined with dashed white lines. mCherry-Pus1 is used as a nuclear marker. The white scale bar represents 3 µm. **F**. Quantification of the percentage of cells containing at least one nuclear GFP-Tel1 focus at each timepoint following release in S phase in the presence of 200 mM HU for WT, *rad53Δ* and *mec1-100* mutant strains. Plotting details as in Figure 1E. **G.** Quantification of the mean number of GFP-Tel1 foci per positive nucleus in each strain. The total number of foci was divided by the number of positive nuclei as in Figure 1F. **H.** Percentage of GFP-Tel1 foci localized in the nucleoplasm relative to foci localized at the nuclear periphery at the indicated timepoints after release in S phase in the presence of 200 mM HU.

Fork reversal has also been described as a pathological state adopted by the fork when facing replication stress in the absence of the effector protein of the DDR Rad53 (Sogo *et al*, 2002; Rossi *et al*, 2015). Of note, the genetic determinants and the outcomes associated to physiological and pathological reversal are unlikely to be similar. If Tel1 presence at the tip is instructed by controlled signaling, pathological situations where the fork structure degenerates because of replication difficulties would not be predicted to elicit GFP-Tel1 foci formation. On the contrary, if Tel1 recruitment to the tip is driven by the mere formation for a DSB-like structure, GFP-Tel1 foci would be expected to form also in these pathological scenarios. To test these hypotheses, we blocked cells lacking Rad53 (*rad53Δ*) in G_1_ and released them synchronously into S phase in the presence of HU, which causes the depletion of dNTP pools. dNTP depletion hampers DNA synthesis, thus creating a replicative stress. For context, in *rad53Δ* cells exposed to HU, more than 10 % of the individual replication intermediates prepared for EM consisted on reversed forks (Sogo *et al*, 2002). In our assay, *rad53Δ* cells manifested a progressive accumulation of GFP-Tel1 foci (Figure 3E) that peaked up at 40 % of the cells (Figure 3F), reaching a mean number of foci per nucleus higher than 2 (Figure 3G). These foci displayed a prevalence (60%) of nucleoplasmic positions (Figure 3H) that was comparable to the proportion observed during physiological fork reversal in WT cells under CPT treatment (65%; Figure 3D).

In the original work by Sogo and colleagues, only the absence of the checkpoint effector Rad53 was tested when assessing its impact on pathological fork reversal, yet authors speculated that it was the lack of a functional checkpoint that gave rise to this situation. To challenge this assumption, we extended the characterization by using a checkpoint-deficient mutant of the upstream DDR sensor kinase Mec1, namely *mec1-100*. This mutant still possesses a proficient kinase activity (Hustedt *et al*, 2015) but is specifically defective for Rad53 activation during replication stress (Paciotti *et al*, 2001).The absence of the checkpoint function of this kinase gave rise to an even sharper appearance of nucleoplasmic GFP-Tel1 foci (Figure 3E), with already about 55% of cells displaying foci, the majority of the latter being nucleoplasmic, only 30 min after S-phase release in HU (Figure 3F). This suggests that, as previously proposed, the lack of a proficient replication checkpoint causes aberrant transitions at stressed replication forks. The two mutants differed at later time points: in *rad53Δ* cells, foci persisted and remained predominantly nucleoplasmic. In *mec1-100* cells, foci became less numerous, and a larger proportion appeared peripheral (Figure 3G,H). This is indicative of the formation of DSBs (Figure 1, (Auboiron *et al*, 2025)), and suggest that, even though replication fork collapse occurs in both *mec1-100* and *rad53Δ* mutants under replication stress (Tercero & Diffley, 2001; Cha & Kleckner, 2002; Rossi *et al*, 2015), it appears faster in the absence of a functional Mec1.

We conclude that GFP-Tel1 very likely marks reversed forks, whether elicited in a regulated or in a pathological manner, by means of the affinity of Tel1 for DNA ends; and that this tool is capable of identifying the transition from reversed to broken forks thanks to the shift from nucleoplasmic to peripheral localization of the detected foci.

### 4. Replication fork slow down caused by deoxyribonucleotide depletion also involves fork reversal in WT cells

In the previous experiment, we had included a control in which WT cells were also challenged with HU to elicit replication stress. Unexpectedly, we observed that up to 20 % of the cells (Figure 3E,F) displayed at least one GFP-Tel1 focus (Figure 3G), which was preferentially localized in the nuclear interior, i.e., in the nucleoplasm (60%; Figure 3H). This observation indicates that replication forks that cannot normally progress due to the lack of dNTPs may frequently engage into fork reversal, a fact that had not been previously observed by EM analyses (Menin *et al*, 2018).

To confirm Tel1 recruitment to replication forks, we asked if, in WT cells, GFP-Tel1 proteins could be found at chromatin positions where replication fork initial stalling occurs in HU. GFP-Tel1 enrichment on chromatin was assessed by <u>Ch</u>romatin Immuno-<u>P</u>recipitation (ChIP) coupled to quantitative PCR (qPCR), and the replication fork position was inferred from the DNA copy number variation calculated from ChIP input qPCR values (Delamarre *et al*, 2020; Forey *et al*, 2021). For ChIP, we used a specific anti-GFP antibody in GFP-Tel1-tagged strains (experimental conditions, Figure 4, in colors) or in an untagged strain (control, Figure 4, in grey). The specific and expected recruitment of Tel1 was validated by monitoring its recruitment to a telomere (*TEL06R*) as a positive control (Figure 4B). We analyzed several genomic positions, some at which replication could be monitored to various extents (early replication origins *ARS306* and *ARS607*), then progressively at longer distances away from them; and some that were devoid of any replication in these experimental conditions (*ARS504* and *ARS305+11,* in which DNA copy number values are 1 or less) (Figure 4A). As a general rule, we observed that, whenever DNA replication could be detected (Figure 4A), a matching, specific GFP-Tel1 signal could be detected, and this in a proportional manner (Figure 4B). This suggested that GFP-Tel1 is recruited to chromatin when forks are stalled by HU treatment. To reinforce this conclusion, we next tested whether DNA copy number correlated with GFP-Tel1 enrichment across the analyzed *loci*. We conducted a proper correlation analysis by asking how variations in variable A (fork position, x-axis) could explain variations in variable B (GFP-Tel1 recruitment, y-axis). The calculated Pearson’s correlation coefficient (*r*) was of 0.68, and of high statistical significance (*p* < 0.0001), meaning that at least 46% (*r^2^ = 0.4624*) of GFP-Tel1 presence of chromatin could be explained (or predicted) by its recruitment at HU-stalled forks (Figure 4C, right). This contrasted with the absolute lack of correlation calculated using the values associated to the unspecific ChIP in the untagged strain (Figure 4C, left). As an additional control, we repeated the experiments in cells where Tel1 cannot be recruited, for the MRX complex protein Rad50 was absent (*rad50Δ*) (Nakada *et al*, 2003). These cells demonstrated Tel1 recruitment levels down to the negative control (Figure 4B).

**Figure 4.**
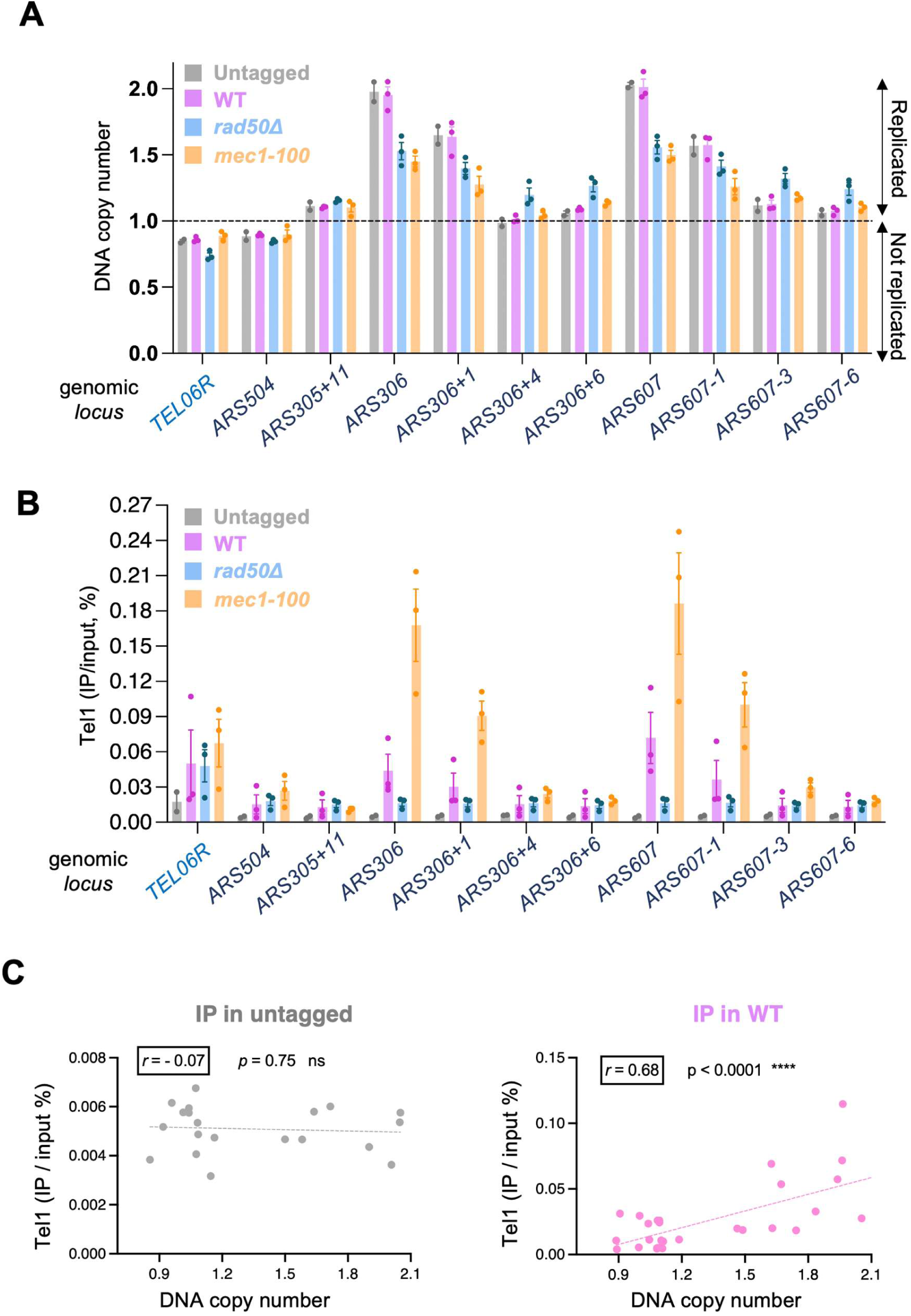
Tel1 is recruited to replication forks during dNTP depletion. **A.** DNA copy number was determined as the ratio between the quantitative PCR (qPCR) values obtained for ChIP input samples of the *loci* of interest and the input of a known unreplicated *locus*, namely *ARS504*. This way, unreplicated *loci* show a ratio close to 1 while replicated *loci* display a ratio between 1 and 2. **B.** ChIP-qPCR analysis of GFP-Tel1 enrichment around early replication origins *ARS306* and *ARS607* in the indicated strains. Cells were released from a G_1_ arrest into S phase for 60 minutes in the presence of 200 mM HU. DNA copy number was determined as the ratio between the quantitative PCR (qPCR) values obtained for ChIP input samples of the loci of interest and the input of two known unreplicated *loci* (mean values), namely *ARS504* and *ARS305*+11. Analyzed *loci* are indicated by their name or by their progressive distance, expressed in kb, from *ARS306* and *ARS607* origins. *TEL06R* was used as a positive control for Tel1 enrichment (Forey *et al*, 2021). Unspecific enrichment corresponds to the qPCR values in an untagged strain (no tag, grey). Mean and SEM correspond to three independent experiments **C.** Correlation analysis between DNA copy number (x-axis) and GFP enrichment (y-axis) at the different analyzed *loci* in the untagged strain. The correlation coefficient (r) and the *p*-value of the analysis are indicated. **D.** Details as in (C), but for the WT strain, in which Tel1 is GFP-tagged.

Last, we observed that Tel1 recruitment at challenged replication forks was much increased in the *mec1-100* mutant (Figure 4B), consistent with the increased proportion of cells containing GFP-Tel1 foci (Figure 3F,G). We conclude that replication fork slowing caused by dNTP depletion involves fork reversal. We propose that forks unable to undertake DNA synthesis because of the dNTP depletion engage in fork reversal, likely transitioning from a replication-competent conformation to a reversed conformation and back, until DNA synthesis can resume. These fork transitions are likely impaired in the absence of a functional replication checkpoint, leading to irreversible fork collapse.

## Discussion

In this work, we report that the kinase Tel1, known for its contribution to different genome integrity preservation pathways, can be tracked by its ability to form nuclear foci whose position informs on its different activities. This way, while peripheral GFP-Tel1 foci mark the early stages of DSB recognition, prior to engagement of its kinase activity, intra-nuclear foci mark reversed replication forks (Figure 5A). However, Tel1’s function at telomeres is not tractable using GFP-Tel1 as a tool. These findings establish a very useful marker to track reversed replication forks and make emerge a new interpretation challenging our view of how replication forks stressed by poor dNTP abundance behave under stress.

**Figure 5.**
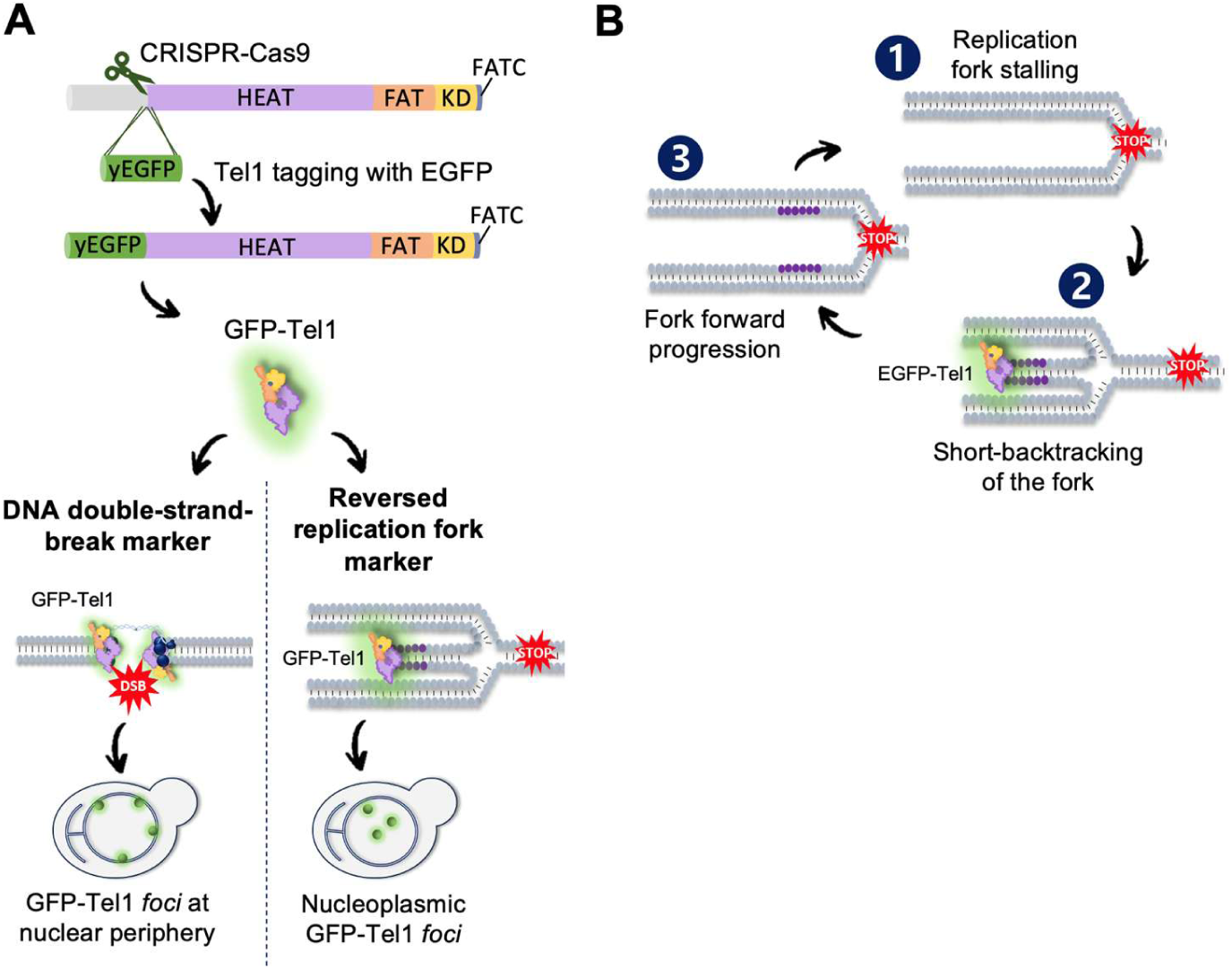
Models. **A.** The Tel1 protein is tagged at its N-terminus with yEGFP using CRISPR-Cas9-mediated genome editing and homologous recombination of a PCR-amplified yEGFP sequence. The resulting strain becomes a tool allowing the discrimination between DSB and reversed forks by analyzing GFP-Tel1 foci’s localization within the nucleus: foci at the nuclear periphery are associated with DSBs, whereas internal, nucleoplasmic foci are associated with reversed replication forks. **B.** Model of back-and-forth replication fork remodeling. We propose that, when cells suffer from (1) replicative stress (i.e., replication fork stalling), it is unlikely that a static configuration is adopted, but that (2) the replication fork undergoes transient reversal that may very dynamically evolve back to (3) a fork configuration competent to resume progression. Multiple cycles of fork reversal and restoration may occur until replication resumption is possible.

### Structural considerations for GFP-Tel1 foci formation

Tel1 recruitment to DNA is made possible by the prior localization to blunt or minimally processed DNA ends by the MRE11-RAD50-NBS1 (MRN complex, MRX in *S. cerevisiae*) (Nakada *et al*, 2003; Falck *et al*, 2005). It has been proposed that, by analogy, Tel1 recruitment to the tip of a reversed fork should be equally dependent on this complex (Menin *et al*, 2018). Our ChIP data suggest that this may be the case, since Tel1 presence at HU-stalled forks fell down to values as those obtained in the control experiment (Figure 4B). Importantly, GFP-Tel1 foci formation in the interior of the nucleoplasm occurred both in regulated, checkpoint-proficient contexts (WT cells exposed to HU or CPT) and in pathological ones (checkpoint-deficient *mec1-100* or *rad53Δ* cells). This suggests that Tel1 recruitment to reversed forks, *via* the MRX complex or not, is a response to the presence of an appropriate DNA substrate and not the outcome of a signaling cascade event.

This interpretation is further supported by the results we have obtained, in response to DSB, using the mutant *tel1-21*, which is incapable of phosphorylating its downstream DDR substrates. In this sense, we asked if Tel1 congregation in the shape of foci necessitates the phosphorylation of its downstream targets, and have found that this is not the case (Figure 1E,F). What is more, likely because Tel1 congregation occurs but reactions do not progress, Tel1 foci accumulate more in the *tel1-21* mutant than in WT cells. Thus, Tel1 concentration on DNA very likely occurs by affinity for the DNA substrate and precedes any signaling event.

Also, as already observed for Tel1 foci formation in response to DNA breaks (Auboiron *et al*, 2025), the mean number of foci observed inside the nucleus in response to fork reversal is low despite many forks potentially being affected (Figure 3). This result suggests that the gathering of multiple forks at a same intra-nuclear position may be necessary in order to reach a threshold allowing enough concentration of Tel1 molecules as to be visualized. Knowing that multiple forks indeed concentrate in space under challenge to enforce adequate fork reversal (González-Acosta *et al*, 2026), this possibility is very likely. The interpretation that Tel1 foci imply the gathering of multiple substrates is further supported by the absence of Tel1 foci detection potentially associated with telomeres (Figure 2). Indeed, while the lack of Tel1 detection associated to short telomeres could be ascribed to a lack of such an event occurring during our experiments, it is very plausible that we failed to detect them because the number of Tel1 molecules congregating at one short telomere are insufficient to elicit the formation of a detectable focus.

An additional structural consideration worth discussing is what drives the formation of peripheral *versus* nucleoplasmic Tel1 foci as a function of the concerned DNA substrate. It could be postulated that the difference in subnuclear localization is dictated by the fact that the DNA breaks under consideration here are two-ended, yet the tip of a reversed fork appears one-ended. However, this is unlikely because, in the *mec1-100* mutant exposed to HU, where replication forks appear to break, the resulting breaks are likely to be one-ended yet they still support the formation of peripheral Tel1 foci. Furthermore, it was already reported that stalled forks tend to remain in the interior of the nucleus (i.e., in WT cells exposed to HU) while those same forks, if forced to break (i.e., in WT cells exposed to HU+ the alkylating agent MMS), relocate to the nuclear periphery (Nagai *et al*, 2008). It is therefore more plausible to invoke that the protein environment surrounding a DSB and a (reversed) fork are drastically different, thus may be able to engage (or not to engage) differently with the nuclear envelope. In this sense, the lipidation of proteins implicated in DNA repair reactions has been shown, in the past, to condition the peripheral *versus* the intra-nuclear position and activity of the concerned protein. As such, Rif1 palmitoylation at residues C466 or C473 anchors the protein to the inner nuclear membrane, thus generating a peripheral compartment favoring DSB repair by non-homologous end joining instead of by homologous recombination (Fontana *et al*, 2019). We put forward the need of assessing how the proteomes of DSB and of reversed forks establish interactions (or not) with the nuclear envelope, and the relevance of its lipidome in this matter. The formation and position of Tel1 foci becomes an advantageous tool to accomplish this task.

### GFP-Tel1 as a novel candidate reporter of fork reversal

In *Saccharomyces cerevisiae*, replication forks in WT cells exposed to HU are not blocked but still progress at a very slow pace (Alvino *et al*, 2007). In contrast, in cultured human cells, fork reversal emerges as a protective, systematic response to HU treatment (Zellweger *et al*, 2015). It has been therefore interpreted that fork reversal in yeast cells is “disfavored”, only being restricted to very specific scenarios (Neelsen & Lopes, 2015). Yet, resection, estimated from the accumulation of RPA at ssDNA by ChIP analysis, was described at HU-stalled replication forks in *S. cerevisiae* (Delamarre *et al*, 2020). In human cells, it is a generally accepted principle that fork reversal precedes fork resection (Mijic *et al*, 2017). This reinforces the notion that fork reversal is a plausible event, in response to HU, also in *S. cerevisiae*. In support, our data reveal that intra-nuclear, GFP-Tel1-labeled events (identical to those marking reversed forks in such specific scenarios) also occur in WT cells exposed to HU. This raises the need for a reconciling explanation. We propose that, in HU, the measurable (even if slow) fork progression would be the positive macroscopic outcome of continuous forward and backward movements of different intensity and duration. As such, fork reversal would be expected to be transient. This may explain why, when assessing fork reversal by electron microscopy in yeast cells, a technique that heavily relies on the frequency and the stability of these DNA intermediates, reversed forks were not detected (Lopes *et al*, 2001). More importantly, it provides an *in vivo* support for the exclusively one-ended DSB-like signals reported in response to WT cells upon HU exposure using sequencing methods (Zhu *et al*, 2019).

Together, our findings suggest a revisited model for the behavior of replication forks in *S. cerevisiae* during HU treatment. We propose continuous rounds of short-lived reversed forks that help stabilize / protect the fork, and that can be subsequently restored into a fork conformation competent for progression, even for short tracts (Figure 5B, model). This would provide a coherent continuum of events that could take place at stressed forks irrespective of the species under consideration. This prompts reevaluation, using nucleoplasmic Tel1 foci as a proxy, of the potential existence of fork reversal in response to genotoxic challenges for which fork reversal was dismissed, such as MMS, H_2_O_2_ or UV light.

Finally, another interesting feature of GFP-Tel1 foci emerged from the localization shift of nucleoplasmic foci to peripheral foci observed in *mec1-100* cells exposed to HU. Thus, following the dynamic localization of GFP-Tel1 foci allows to track replication fork processing, from reversal to collapse. Interestingly, this shift of localization was not observed in the *rad53Δ* mutant in HU, in which GFP-Tel1 foci persisted being nucleoplasmic. It is assumed that forks collapse occurs in HU whenever the checkpoint is not functional, yet here our tool allowed to uncover a major difference that could challenge this assumption. Our data suggest that, while the mutant Mec1-100 may not be able to support any kind of phosphorylation dynamics at challenged forks, thus leading to their collapse and emergence of DSBs, the absence of Rad53 does not preclude wild-type Mec1 protein from still launching protective strategies at forks, that is, preventing fork collapse and immediate DSB generation. Revising the literature shows that previous evidence of DNA breakage in cells lacking Rad53 in the presence of HU did not manifest before three hours of treatment, with cells already reaching late S phase, as shown by cytometry (Rossi *et al*, 2015). This strongly points at these *rad53Δ* cells suffering from processing of incompletely replicated DNA tracks by nucleases in late S/G_2_, rather than challenged forks that break on-site. Our results are therefore consistent with previous observations (Cha & Kleckner, 2002) and with the separate roles of Mec1 and Rad53 checkpoint kinases in stabilizing replication forks in the face of other genotoxic stresses (Segurado & Diffley, 2008).

### On the role of Tel1 at forks: considerations from different models

In response to CPT, reversed forks were detected in *S. cerevisiae* to which Tel1 was proposed to be recruited (Menin *et al*, 2018). We indeed detected GFP-Tel1 nucleoplasmic foci under these conditions (Figure 3B-D). At these forks, Tel1 was proposed to restrict unscheduled nucleolytic activities by Mre11, implying Tel1’s role at these structures would be of a protective nature. An overview of the current literature asking if ATM, the Tel1 homolog in human cells, fulfils a similar role at reversed forks, leads to a somewhat contradictory idea.

By applying systematic proteomics profiling of replication forks, ATM has been reported to physically relocate to replication forks once they break, which was achieved using high doses (1 µM) of CPT. On the contrary, whenever DNA replication was slowed down using HU, and likely forks were reversed, ATM was not detected at such challenged replication forks (Nakamura *et al*, 2021). This therefore supports the canonical view of ATM^Tel1^ locating, exclusively, at *bona fide* breaks, and not at structures such as the tip of a reverse fork. Beyond, ATM activities in authorizing fork reversal have been reported, although these do not demonstrate the physical presence of ATM at the reversed fork itself. For example, the TIP60-ATM pathway supports the recruitment of the reverse-inducing factor SMARCAL1 to stalled forks (Schleicher *et al*, 2022). The tip of the reversed fork is said to attract and stabilize the MRN complex, which itself is known to attract ATM, which in turn imparts a “priming” phosphorylation onto MRN that is important to regulate downstream tip processing by resection and concomitant fork stability (Zhang *et al*, 2025). In *Xenopus* extracts, ATM was found to be activated concomitantly with fork reversal, which in turn engage resection of the reversed fork intermediate. The study claims reversed forks as substrates for ATM activation and reveals a phospho-regulatory circuit that governs reversed fork processing during inter-strands crosslink repair” (Altshuller *et al*, Preprint). A recent pre-print highlights a dominant effect exerted by ATM hyper-signaling at reversed forks that necessitates to be counteracted by the phosphatase PPM1D in order to enable proper fork restart (Cao *et al*, Preprint). Last, cyclic GMP-AMP synthase (cGAS), a DNA sensor activating type-I interferon response, can bind to chromatin and slow down DNA replication forks unless countered by ATM, as ATM promotes its disengagement from chromatin (Song *et al*, 2026). Thus, it remains contradictory whether Tel1 and ATM protect forks from resection, or favors it.

### Collateral observations: learning more

The use of GFP-Tel1 foci as a tool to identify DSB- and reversed fork-associated phenomena fosters new discoveries. For example, initially described in human cells, the CST complex was reported to associate with DNA Polymerase α/primase to fill-in ssDNA gaps, thus limiting 3’ overhangs (Mirman *et al*, 2018, 2022). This sparked a deeper conception of the role(s) of the CST complex beyond telomeres (Lyu *et al*, 2021). Recently, similar findings were reported in *S. cerevisiae* (Ilioaia *et al*, 2025). Our data show that the CST mutant *cdc13-1*, at the restrictive temperature, forms many more robust GFP-Tel1 foci in response to zeocin, reinforcing the idea that the CST complex plays a role in DNA repair also in *S. cerevisiae*. However, since the absence of the CST complex in response to DSBs will be accompanied by extensively resected DNA, it is difficult to reconcile such long ssDNA 3’ end overhangs with more Tel1-marked events. This apparent paradox deserves further exploration.

Last, it is worth commenting that frequent, bright GFP-Tel1-positive structures were observed in the cytoplasm of the *cdc13-1* mutant when cultured at the restrictive temperature (Figure 2C). These signals are reminiscent to the ones we reported when forcing cytoplasmic Tel1 retention in the cytoplasm. This was achieved by increasing the pool of the lipid PI4P at the Golgi apparatus (by ablation of its phosphatase Sac1), which we have shown can titrate Tel1/ATM away from the nucleus, both in S*. cerevisiae* and in human cells (Ovejero *et al*, 2023). Crucially, an independent study reports that the Golgi protein Vps74, which promotes this same PI4P dephosphorylation step, helps maintain low levels of the CST protein Stn1 (Rodrigues *et al*, 2018). First, we infer that PI4P consumption by dephosphorylation is important to maintain low levels of Stn1. Second, the increase in GFP-Tel1 cytoplasmic structures upon Sac1 ablation (Ovejero *et al*, 2023) or CST deficiency (Figure 2C) (both events related to the Golgi pools of PI4P) warrant further assessment of a potential Golgi-mastered control of telomere homeostasis.

## Material and Methods

*Reagents* used in this work for cell treatments were zeocin (R25001, ThermoFisher Scientific); camptothecin (CPT, C9911 Sigma-Aldrich); hydroxyurea (HU, H8627 Sigma-Aldrich); and α-factor, synthesized upon demand by BIOTEM.

### Strains, plasmids and oligonucleotides

The strains used in this study are presented in Table 1, the plasmids in Table 2 and oligonucleotides in Table 3. Strains were obtained either by classical methods for integration, transformation and crosses, or by CRISPR-Cas9 technology in the case of Tel1 yEGFP-tagging (DiCarlo *et al*, 2013) as previously described by us (Auboiron *et al*, 2025). *tel1-21* mutant was created as described by Ogi *et al*. by mutating Ile2779 and Tyr2780 both into Ala by CRISPR-Cas9, then tagged with yEGFP also by CRISPR-Cas9. All strains were transformed with linearized mCherry-Pus1 plasmid digested by *Bgl*II restriction enzyme (R0144, New England Biolabs) to visualize nuclei in microscopy experiments. The *cdc13-1* mutant was kindly provided by Brian Luke, then tagged for EGFP-Tel1 and mCherry-Pus1, as the other strains in this study.

**Table 1.** Strains used in this study.

| Simplified Genotype | Full genotype | Source |
|---|---|---|
| WT, yEGFP-Tel1 | MAT a, *ade2-1, his3-11,15, can1-100, leu2-3,112, trp-11, ura3-1, GAL+, psi+, RAD5, yEGFP-Tel1* | PP3952 (Forey *et al*, 2021) |
| WT, yEGFP-Tel1 mCherry-Pus1 | MAT a, *ade2-1, his3-11,15, can1-100, leu2-3,112, trp-11, ura3-1, GAL+, psi+, RAD5, yEGFP-Tel1, mCherry-Pus1::URA3* | MM-40 (Auboiron *et al*, 2025) |
| WT, yEGFP-Tel1, mCherry-Pus1 (for CPT treatment) | MAT a, *ade2-1, his3-11,15, can1-100, leu2-3,112, trp-11, ura3-1, GAL+, psi+, RAD5, yEGFP-Tel1, mCherry-Pus1::URA3, POL31-AID\*-6FLAG::HPH<sup>R</sup>, OsTIR-9myc::ADE2, yEGFP-Tel1, mCherry-Pus1::URA3* | MM-611 - This study |
| *mrc1Δ*, yEGFP-Tel1 mCherry-Pus1 (for CPT treatment) | MAT a, *ade2-1, his3-11,15, can1-100, leu2-3,112, trp-11, ura3-1, GAL+, psi+, RAD5, yEGFP-Tel1, mCherry-Pus1::URA3, POL31-AID\*-6FLAG::HPH<sup>R</sup>, OsTIR-9myc::ADE2, yEGFP-Tel1, mCherry-Pus1::URA3 mrc1Δ::TRP1* | MM-989 - This study |
| *yEGFP-tel1-21*, mCherry-Pus1 | MAT a, *ade2-1, his3-11,15, can1-100, leu2-3,112, trp-11, ura3-1, GAL+, psi+, RAD5, yEGFP-tel1-21 (I2779A, Y2780A), mCherry-PUS1::URA3* | MM-1200 and MM-1201 - This study |
| *cdc13-1*, yEGFP-Tel1, mCherry-Pus1 | MATa, *met15Δ0, his3Δ1, leu2Δ0, ura3Δ0, cdc13::cdc13-1::NAT<sup>R</sup>, yEGFP-Tel1, mCherry-Pus1::URA3* | MM-1161 and MM-1162 - This study (Modified from Brian Luke lab) |
| *rad53Δ, sml1Δ*, yEGFP-Tel1, mCherry-Pus1 | MAT a, *ade2-1, his3-11,15, can1-100, leu2-3,112, trp-11, ura3-1, GAL+, psi+, RAD5, yEGFP-Tel1, mCherry-PUS1::URA3, rad53Δ::KAN<sup>R</sup>, sml1Δ::HIS3* | MM-1157 and MM-1158 - This study |
| *mec1-100*, yEGFP-Tel1, mCherry-Pus1 | MATa, *ade2-1, trp1-1, can1-100, leu2-3,112, his3-11,15, ura3, GAL, psi+, RAD5, yEGFP-Tel1, mec1-100::LEU2::mec1::TRP1, mCherry-Pus1::URA3* | MM-1369 and MM1370 - This study |
| *mec1-100*, yEGFP-Tel1 | MATa, *ade2-1, trp1-1, can1-100, leu2-3,112, his3-11,15, ura3, GAL, psi+, RAD5, yEGFP-Tel1, mec1-100::LEU2::mec1::TRP1* | PP5181 - This study |
| *rad50Δ*, yEGFP-Tel1 | MATa, *ade2-1, trp1-1, can1-100, leu2-3,112, his3-11,15, ura3, GAL, psi+, RAD5, yEGFP-Tel1, rad50Δ::KAN<sup>R</sup>* | PP5218 - This study |

**Table 2.** Plasmids used in this study.

| **Simplified Name** | **Detailed Information** | **Source** |
|---|---|---|
| pMEL10-Tel1p | pMEL10-gRNA(*TEL1p*) | (Auboiron *et al*, 2025) |
| p-*TEF1p-CAS9* | pRS414-*TEF1p-CAS9-CYC1t* | (DiCarlo *et al*, 2013) Addgene #43802 |
| p-yEGFP | pKT128 | (Sheff & Thorn, 2004) |
| p-mCherry-*PUS1* | YIplac211-mCherry-*PUS1* | (Han *et al*, 2008) |

**Table 3.**
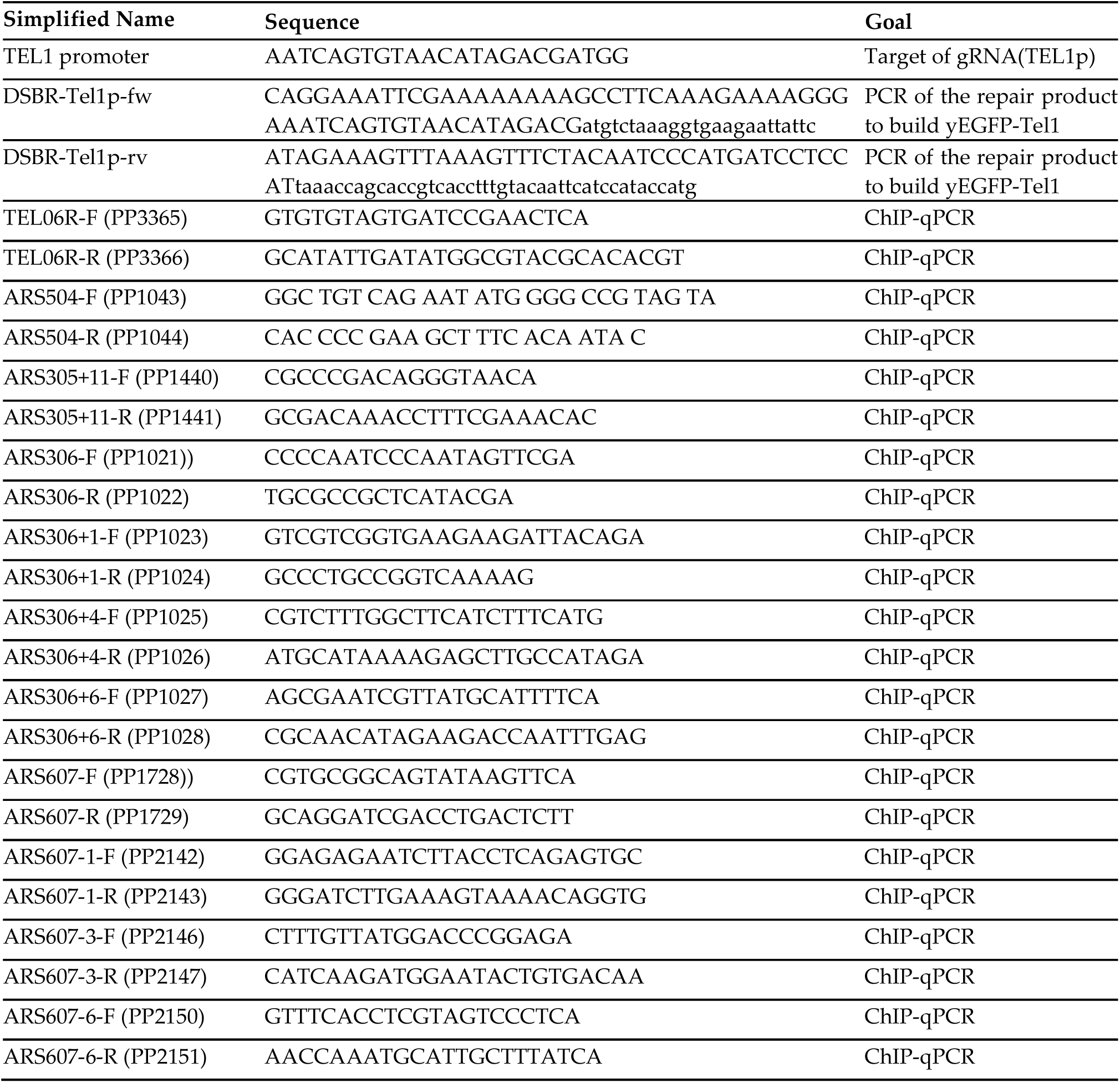
Relevant sequences and primers used in this study.

### Culture and treatments

*Saccharomyces cerevisiae* cells were grown in YEP + 2% glucose for all experiments. DSB are induced by zeocin addition (100µg/mL final concentration) after the cells reach exponential growth. Cells were collected for microscopy every 30min for 210min. To accomplish experiments with the *cdc13-1* mutant, cultures were firstly incubated at the permissive temperature (23°C) and split when they reached exponential growth, cells were then incubated at the permissive (23°C) or the restrictive (37°C) temperature with or without zeocin addition (100µg/mL final concentration). Cells were collected for microscopy every hour for 3h. For CPT treatment experiments, cells were treated with 100 µM CPT and harvested after 90 and 180 min. To synchronize cells, three shots of α-factor (each dose at 2µg/mL final concentration) were added to the medium to synchronize cells in G_1_ – one dose at the starting time point, two doses after 1h of incubation and one last dose after 2h of incubation for a total of 3h. After G_1_ synchronization, cells were centrifuged, washed 2 times with water to release them in S phase, either with or without HU (0.2M final concentration) in fresh YEP + 2% glucose. Cells were collected for microscopy every 30min for 210min, or after 60min for ChIP experiments.

### Western-Blotting

For GFP-Tel1 detection, approximately 1.10^8^ exponentially growing cells were collected and washed with 20% trichloroacetic acid (TCA) at 4°C. Cell pellets were resuspended in 200 µL of ice-cold 20% TCA, and ∼200 µL of glass beads were added to disrupt cells by vortexing for 10 min at 20-25°C. After centrifugation, the pellet was resuspended in 100 µL of 2× Laemmli sample buffer which turns yellow. The minimum volume of 1 M Tris base (without pH adjustment) was added to restore the blue color. The final volume was adjusted to 200 µL with water. Samples were boiled for 10 min at 96°C and clarified by centrifugation. 25 µl microliters of each protein extract were loaded onto a 3–8% precast Tris-acetate gradient gel (Bio-Rad) and separated by electrophoresis for 75 min at 150 V in Tris-acetate running buffer. Proteins were transferred onto nitrocellulose membranes in Tris-glycine transfer buffer supplemented with 20% methanol at 30 V for 16 h at 4°C. Membranes were then stained with Ponceau S to verify protein transfer. Immunoblotting was performed using a rabbit anti-GFP primary antibody (1:2000; Clinisciences, TP-401) overnight at 4°C, followed by incubation with a fluorescent anti-rabbit secondary antibody (1:15000; Fisher Scientific, 10733944) for 30 min at 20-25°C. Fluorescent signals were detected by an Odyssey Scanner M (LI-COR).

### Fluorescence Microscopy

1 mL of the culture of interest was centrifuged, the supernatant thrown away and the pellet resuspended in the remaining 50 μL. 3 μL of this living-cell suspension were directly mounted on a microscope slide for immediate imaging of the pertinent fluorophore-tagged protein signals. Imaging was achieved using a Zeiss Axioimager Z2 microscope with an objective 63X Plan Apochromat 1.4 NA oil. Visualization, co-localization and analysis were done using Image J.

### Chromatin immunoprecipitation (ChIP) and quantitative PCR

5.10^8^ exponentially growing cells in YPAD medium (50 mL at 1.10^7^ cells/mL) were blocked in G_1_ by the addition of 100 µg of α-factor peptide (GenScript, RP01002) every 45 min for 135 min at 25°C. Cells were released from the G_1_ arrest by filtration and resuspension in fresh medium containing 200 mM Hydroxyurea (Cliniscience, Cat#AG000WIJ) and collected 60 min later. Cells were crosslinked for 15 min with 1% formaldehyde (Sigma F8775) at 25°C under agitation. Fixation was quenched by addition of 250 mM Glycine (Sigma G8898) for 5 min under agitation. Cells were washed two times by centrifugation with cold TBS (4°C). Pellets were frozen and conserved at -20°C until processed. Cell pellets were resuspended in 1.2 mL of Lysis buffer (50 mM HEPES-KOH pH7.5, 140 mM NaCl, 1 mM EDTA, 1% Triton X-100, 0.1% Na-deoxycholate) supplemented with 1 mM PMSF and anti-protease cocktail (cOmplete Tablet, Roche, 505649001) and lysed by beads-beat method (MB400 U, Yasui Kikai, Osaka). Recovered lysates adjusted to 3 mL with Lysis buffer were sonicated with a Q500 sonicator (Qsonica) for 12 cycles (15 s ON, 45 s OFF, amplitude 50) and cleared by centrifugation for 4 min at 2,400 *g*. Aliquots (Input samples) of the resulting whole cell extracts (WCE) were kept for quantitative PCR (qPCR, 135 µL), DNA fragmentation control by agarose gel electrophoresis (120 µL) and immunoprecipitation (IP) control by western blot (25 µL). For the IP, 120 µL of Dynabeads Prot A (Life Technologies, Cat#100.02D) were washed three times and resuspended in 1 mL of PBS, 0.5% BSA, 0.1% Tween and incubated with 10 µL of anti-GFP antibody (Thermo Fisher Scientific, Cat#A-11122) on a rotating wheel for six hours at 4°C. Antibody-coupled Dynabeads were washed three times with 1 mL of PBS, 0.5% BSA, 0.1% Tween and added to the remaining 2.7 mL of WCE, then incubated overnight on a rotating wheel at 4°C. After immunoprecipitation, beads were immobilized on a magnetic rack. 25 µL of the supernatant were collected for western blot (Flow-through sample) and beads were washed on ice with cold solutions: twice with Lysis buffer (50 mM HEPES-KOH pH7.5, 140 mM NaCl, 1 mM EDTA, 1% Triton X-100, 0.1% Na-deoxycholate), twice with Lysis buffer + 0.36 M NaCl (50 mM HEPES-KOH pH7.5, 360 mM NaCl, 1 mM EDTA, 1% Triton X-100, 0.1% Na-deoxycholate), twice with Wash buffer (10 mM Tris-HCl pH8, 0.25 M LiCl, 0.5% IGEPAL, 1 mM EDTA, 0.1% Na-deoxycholate) and once with TE buffer (10 mM Tris-HCl pH8, 1 mM EDTA). Immunoprecipitated proteins were uncoupled from beads by incubation for 11 min at 65°C with 140 µL of Elution Buffer (50 mM Tris-HCl pH8, 10 mM EDTA, 1% SDS) with occasional shaking. 5 µL of the eluate were collected for western blot (IP sample) and the remaining 135 µL were incubated alongside with the Input samples at 65°C overnight after the addition of 135 µL of TE buffer +0.1% SDS for de-crosslinking. 130 µL of TE containing 60 mg RNase A (Sigma, R65-13) were added to the samples and incubated for 2 hours at 37°C. Proteins were digested by addition of 20 µL of 20 mg/mL Proteinase K (Sigma, P6556) and incubated for 2 hours at 37°C. 50 µL of 5 M LiCl were added to the samples before DNA purification with two extraction rounds using equal volumes of Phenol: Chloroform: Isoamyl Alcohol 25:24:1 (Sigma, P2069). The aqueous phase containing the DNA was precipitated by addition of 100 mM Sodium Acetate (Sigma, S2889), 26 mg/ml of Glycogen (Roche, 10901393001) and 2.5 volumes of ice-cold ethanol, thoroughly mixed and incubated overnight at -20°C. Samples were centrifuged for 45 min at 21,000 *g* at 4°C. DNA pellets were washed with ice-cold 70% ethanol, air-dried and resuspended in H20. qPCR reactions were performed with SYBR® Green (Roche) and LightCycler480 (Roche) on 384 well-plates. The program consisted on one cycle of heat activation (95°C, 10min), 45 cycles of PCR cycling (95°C – 15sec; 60°C – 15 s; 72°C – 15 s) and one cycle of melting curves (from 60°C to 95°C). Primer sequences are indicated in Table 3. IP/Input ratios were calculated and plotted using GraphPad Prism 10. An unspecific IP was performed with an untagged Tel1 strain and the IP efficiency was controlled by SDS-PAGE after addition of equal volumes of 2X Laemmli buffer to the Input, Flow-through and IP samples and boiled at 95°C for 30 min.

### Quantifications, Graphical Representations and Statistical Analyses

The number of nuclei displaying foci of the analyzed proteins as well as the number of foci present per nucleus were counted visually by the experimenter. To determine whether GFP-Tel1 foci were localized at the nuclear periphery or within the nucleoplasm, the mCherry-Pus1 signal was used as a nuclear marker. Foci were systematically classified by the experimenter as peripheral when strictly located at the border of the mCherry-Pus1 signal and as nucleoplasmic when located away from such border. The details and the rigor of this strategy, challenged by using a nucleoporin as a solid reference, have been already reported in our previous work (Auboiron *et al*, 2025). GraphPad Prism was used both to plot the graphs and to statistically analyze the data. The mean value of nuclei displaying foci was calculated for each independent experiment, and the SEM (standard error of the mean) was used to inform on inter-experiment variation. The SEM estimates how far the calculated mean is from the real mean of the sample population, while the SD (standard deviation) informs about the dispersion (variability) of the individual values constituting the population from which the mean was drawn. Given that the goal of our error bars was to describe the uncertainty of the true population mean being represented by the sample mean, we chose to plot the SEM. To plot frequencies, we divided the actual observations from each category (number of foci per nucleus) by the total number of observations.

## Author Contributions

Conceptualization: M.A., B.P., M.M.-C.; Data curation: M.A., K.S., A.B., B.P., M.M.-C.; Formal Analysis: M.A., K.S., A.B., B.P., M.M.-C.; Funding acquisition: P.P., M.M.-C.; Investigation: M.A., K.S., A.B., M.M.C.; Methodology: M.A., K.S., A.B., B.P., M.M.-C.; Project administration: M.M.-C.; Supervision: B.P., M.M.-C.; Visualization: M.A., M.M.-C; Writing – original draft: M.A., B.P., M.M.-C.; Writing – review & editing: M.A., K.S., A.B., P.P., B.P., M.M.-C.

## Acknowledgments

We thank Brian Luke for sharing the mutant *cdc13-1*. We are thankful to the joint IGMM-CRBM ‘yeast media and technologies service’ for providing us with ready-to-use media. We acknowledge the imaging facility MRI, member of the national infrastructure France-BioImaging, supported by the French National Research Agency (ANR-24-INBS-0005 FBI BIOGEN). We thank the French National Research Agency (ANR) for supporting this work *via* two different grants: “Genome Sterolity” (ANR-21-CE12-0004-01) and “SELDOM” (ANR-24-CE11-3401-02).

## Abbreviations

cGAS: cyclic GMP-AMP synthase
ChIP: Chromatin Immunoprecipitation
CPT: camptothecin
CST: Cdc13-Stn1-Ten1 complex
DDR: DNA damage response
DSB: DNA double strand break
EGFP: enhanced green fluorescent protein
HU: hydroxyurea
MRN: Mre11-Rad50-Nbs1 complex
PIKK: phosphatidyl-inositol-3-kinase-related kinase
SEM: standard error of the mean
WCE: whole cell extract
WT: wild type.

